# Application of Transfer Learning for Rapid Calibration of Spatially-resolved Diffuse Reflectance Probes for Extraction of Tissue Optical Properties

**DOI:** 10.1101/2023.10.23.563629

**Authors:** Md Nafiz Hannan, Timothy M. Baran

## Abstract

**Significance:** Treatment planning for light-based therapies including photodynamic therapy requires tissue optical property knowledge. These are recoverable with spatially-resolved diffuse reflectance spectroscopy (DRS), but requires precise source-detector separation (SDS) determination and time-consuming simulations.

**Aim:** An artificial neural network (ANN) to map from DRS at short SDS to optical properties was created. This trained ANN was adapted to fiber-optic probes with varying SDS using transfer learning.

**Approach:** An ANN mapping from measurements to Monte Carlo simulation to optical properties was created with one fiber-optic probe. A second probe with different SDS was used for transfer learning algorithm creation. Data from a third were used to test this algorithm.

**Results:** The initial ANN recovered absorber concentration with RMSE=0.29 µM (7.5% mean error) and µ_s_^’^ at 665 nm (µ_s,665_^’^) with RMSE=0.77 cm^-1^ (2.5% mean error). For probe-2, transfer learning significantly improved absorber concentration (0.38 vs. 1.67 µM, p=0.0005) and µ_s,665_^’^ (0.71 vs. 1.8 cm^-1^, p=0.0005) recovery. A third probe also showed improved absorber (0.7 vs. 4.1 µM, p<0.0001) and µ_s,665_^’^ (1.68 vs. 2.08 cm^-1^, p=0.2) recovery.

**Conclusions:** A data-driven approach to optical property extraction can be used to rapidly calibrate new fiber-optic probes with varying SDS, with as few as three calibration spectra.

## 1 Introduction

Photodynamic therapy (PDT), which produces an antimicrobial effect through the photochemical production of reactive oxygen species, is a promising treatment modality for eradication of infectious disease[1]. We are particularly interested in the use of PDT to treat deep tissue abscesses, which are localized collections of infected, purulent fluid surrounded by a fibrous pseudo-capsule. We have shown that PDT is safe and feasible for this purpose in a Phase 1 clinical trial[2], and that PDT is efficacious against bacteria cultured from human abscesses[3]. Effective PDT requires sufficient doses of both the photosensitizer and treatment light. As the light dose is dependent upon the optical properties of the target tissue, namely absorption and scattering, accurate measurement of tissue optical properties is required for designing efficacious PDT treatment plans. To that end, we have constructed and validated a spatially-resolved diffuse reflectance spectroscopy system that is capable of extracting tissue optical properties using a fiber-optic probe placed in contact with the abscess wall[4]. Critically, the fiber-optic probe used to deliver light and collect diffusely reflected light was required to fit through the drainage catheter used for standard of care abscess drainage, which has an inner diameter of only 2 mm. This spectroscopy system was utilized during our recently completed Phase 1 clinical trial[2] to quantify human abscess wall optical properties [5]. Substantial inter-patient variability in the abscess wall optical properties was found[5]. We have shown that patient-specific treatment planning incorporating knowledge of these optical properties greatly improves delivery of an efficacious fluence rate to the abscess wall[6, 7]. Measurement of individual patient optical properties is therefore of high importance for the clinical success of PDT for this application.

To extract optical properties from diffuse reflectance spectroscopy, different approaches have been used, broadly categorized into two groups: analytical and numerical. Analytical approximations to the radiative transport equation such as the diffusion[8, 9] and P_3_ [10] approximations have been utilized by many research groups. However, these approximations place limitations on source-detector separation, requiring detectors to be multiple transport mean free paths from the source. As described above, our primary clinical target requires that a fiber optic probe with multiple source detectors separations (SDS) must fit through a catheter with a diameter of 2 mm. As a consequence, these approximations are not valid for the probes we aim to deploy clinically, and more broadly for spatially-resolved diffuse reflectance at short SDS. Many research groups have therefore employed numerical approaches, such as the Monte Carlo lookup table (MCLUT) inverse model for optical property recovery[4, 11]. This approach is more flexible in terms of SDS range and is also valid for a wide range of optical properties. However, the usage of MC requires accurate SDS measurements for model creation, and generation of an extensive MCLUT which remains resource and time-intensive despite graphics processing unit (GPU) acceleration. Moreover, a distinct MC library must be generated for every probe employed. This is undesirable for cases where similar but distinct probes need be deployed, such as for single-use devices.

With advancements in computational power and machine learning (ML) algorithms, numerous research groups have turned to fully data-driven ML approaches for recovering optical properties from diffuse reflectance spectroscopy (DRS). Farrell *et al* pioneered the use of an artificial neural network (ANN) trained with the diffusion approximation to extract optical properties from simulated experimental data[8]. Recognizing the limitations of the diffusion approximation, Kienle *et al* subsequently employed an ANN trained with MC simulations[12]. More recently, Nguyen *et al* demonstrated that deep learning models showed lower error and faster runtime compared to a MCLUT inverse model[13]. Lan *et al* developed an ANN model that achieved comparable accuracy to the traditional Monte Carlo-based inverse model while offering improved speed and flexibility[14].

However, these prior studies were confined to the performance of ANN-based optical property recovery for a single device. Data obtained from analytical solutions [8] and/or MC libraries [12, 14] for a specific probe in combination with phantom data collected by that probe were used to train these ANN models. None of these studies showed extension of ANNs to the case of unseen probes with variable SDS. As the ANNs created by these groups were probe-specific, they are not suitable for clinical adoption or more widespread application where rapid calibration of a large number of new devices is required. An ANN created for one specific device cannot be expected to perform well for other probes having dissimilar SDS, as the DRS signal will vary in unknown ways. To circumvent this with current methods, either the probe SDS would need to be very tightly controlled or a separate ANN would need to be created for each probe. Both of these solutions would impede clinical implementation by increasing costs due to tight tolerancing and prolonging the time required for calibration.

To facilitate our envisioned multi-center clinical trials and eventual widespread clinical adaptation, we propose a Transfer Learning (TL) approach for rapid probe-to-probe calibration. TL is a widely employed approach in the ML field, wherein models trained for one specific task are partially retrained to address a related task that has limited available data. We hypothesize that by using a feature-extraction based TL algorithm, an ANN created for one probe can be adapted to similar probes via a small dataset consisting of only a few calibration spectra collected by the target probe. This will enable the creation of an initial ANN for a single probe, which can be easily adapted to target probes without the need for the time and resource-intensive steps mentioned earlier, thus rendering large-scale clinical adoption more feasible.

This study followed a three-step workflow. First, an initial artificial neural network (ANN) was created and validated using data captured with a single fiber optic probe (probe-1). This initial ANN served as a pre-trained model for subsequent transfer learning steps. Second, we utilized data from a second fiber optic probe with slight differences in source-detector separation (probe-2) to create and validate a TL algorithm, which included selecting and optimizing the TL algorithm, TL dataset size, and associated hyperparameters. Finally, this TL algorithm was applied to data from a third fiber optic probe (probe-3) in a “real-world” scenario to independently assess its performance and applicability.

## 2 Methods

### 2.1 Spectroscopy System

The fiber optic probe-based diffuse reflectance system utilized in this study has been extensively described in our earlier publication [4]. Briefly, it is a continuous wave spectroscopy system in which broadband light from a tungsten halogen lamp (HL-2000-HP-FHSA, Ocean Optics, Inc., Largo, FL) is emitted from a source fiber into the sample. Eight detector fibers, which are placed at different source-detector separations (SDS) from the source fiber on the face of the probe, collect the reflected light for detection by a spectrometer (QE Pro, Ocean Optics). An encrypted, password-protected laptop computer is used to operate this system via a custom interface created in LabVIEW (National Instruments, Austin, TX).

### 2.2 Probe Characterization

Three custom-made fiber optic probes were used, each having an outer diameter of 2 mm. These probes were manufactured by Pioneer Optics Company (Bloomfield, CT), following the same design specifications. These specifications involved positioning the fiber faces at the distal end of the probe in a “plus” pattern, with a desired distance of 300 μm between adjacent fiber centers. While the design intended for each of these probes to possess the same SDS, the delivered probes had varying SDS values compared to each other. At the short separations used in these probes, precise knowledge of SDS is important for accurate recovery of tissue optical properties.

In order to determine the actual SDS for each probe, a stereomicroscope (SMZ1500, Nikon Instruments, Melville, NY) was used. The distal end of each probe was imaged both with and without a US Air Force resolution target (USAF 1951 1X, Edmund Optics, Inc., Barrington, NJ) in the same frame. The air force target was used as a distance reference. The centers of each fiber were identified and SDS for each fiber were determined. An image of the distal face of probe-1 showing the position of the source and detector fibers is presented in Figure 1. The SDS for all 8 detectors fibers for each probe are given in Table 1.

**Figure 1:**
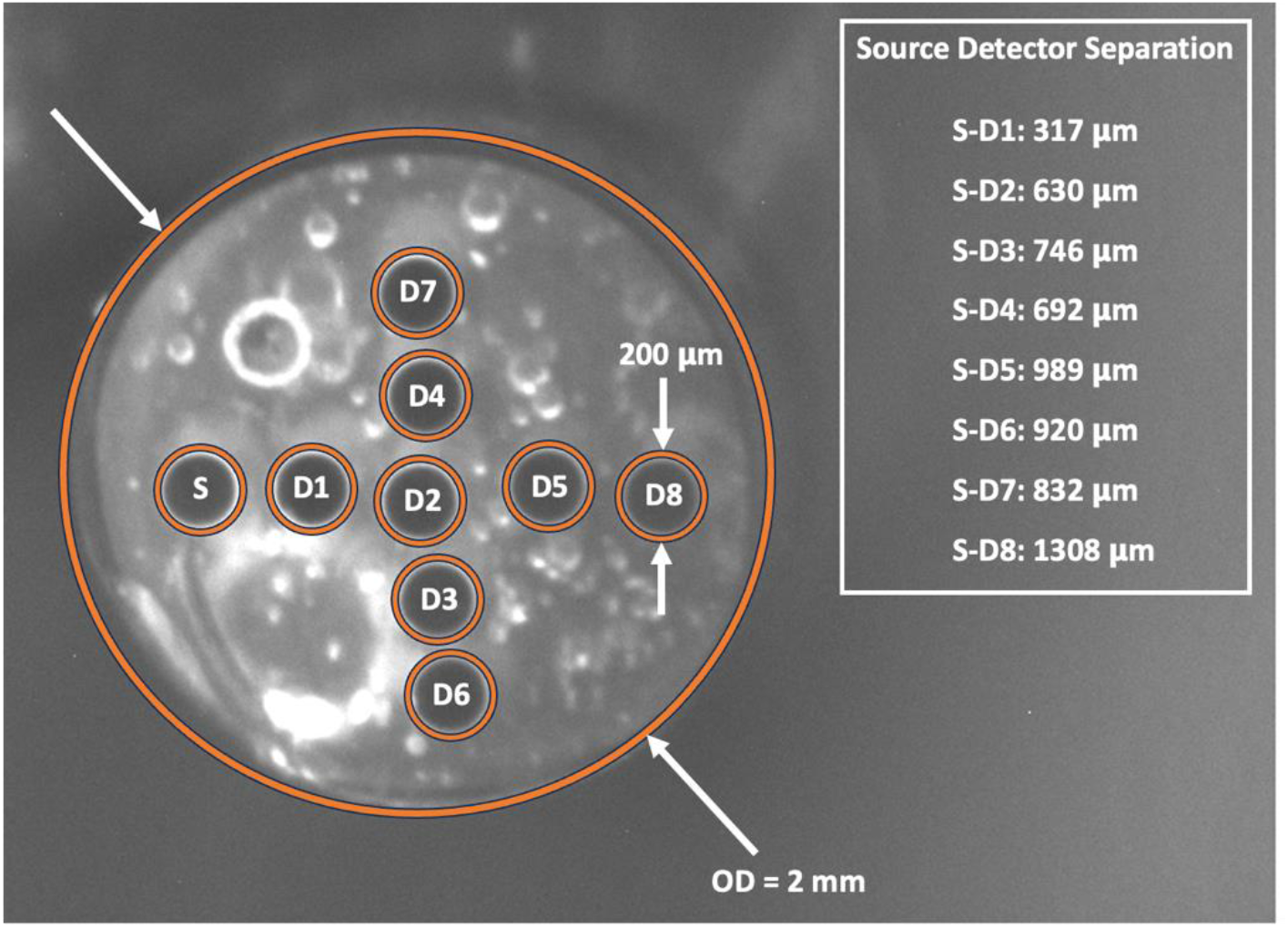
Image of the distal face of probe-1 showing the 200 µm diameter fibers. The source fiber (S) and detector fibers (D1-D8) are denoted using orange circles. The outside diameter is abbreviated as “OD.”

**Table 1:** Source detector separation in µm for the 8 detector fibers of probes 1, 2 and 3.

| Probe No. | Detector Fiber 1 | Detector Fiber 2 | Detector Fiber 3 | Detector Fiber 4 | Detector Fiber 5 | Detector Fiber 6 | Detector Fiber 7 | Detector Fiber 8 |
| --- | --- | --- | --- | --- | --- | --- | --- | --- |
| Probe 1 | 317 | 630 | 746 | 692 | 989 | 920 | 832 | 1308 |
| Probe 2 | 315 | 648 | 738 | 761 | 985 | 914 | 911 | 1342 |
| Probe 3 | 325 | 736 | 804 | 829 | 1082 | 951 | 990 | 1424 |

### 2.3 Artificial Neural Networks

#### 2.3.1 Creation and Validation of Initial ANN (ANN_1_)

An artificial neural network (ANN) approach was employed to recover optical properties from diffuse reflectance spectra (DRS). An initial ANN (ANN_1_) was trained to map from the experimentally collected DRS for probe-1 to corresponding optical properties (absorption and reduced scattering spectra). It had 8 input nodes, one for each SDS, and 2 output nodes, one for each optical property (absorption and reduced scattering). It was created by combining two separate ANN ensembles, illustrated in Figure 2. The first ANN ensemble (ANN_EXP1-MC1_) takes in experimental diffuse reflectance intensities at the 8 detector fibers as inputs, with outputs corresponding to Monte Carlo simulated reflectance intensities for these 8 fibers. The second ANN ensemble (ANN_MC1-OP_) takes in these 8 simulated reflectance intensities and outputs corresponding optical properties. To get the final output of the ensemble models, predictions of individual ANNs were averaged. ANNs used in this research were created using the Keras API [15].

**Figure 2:**
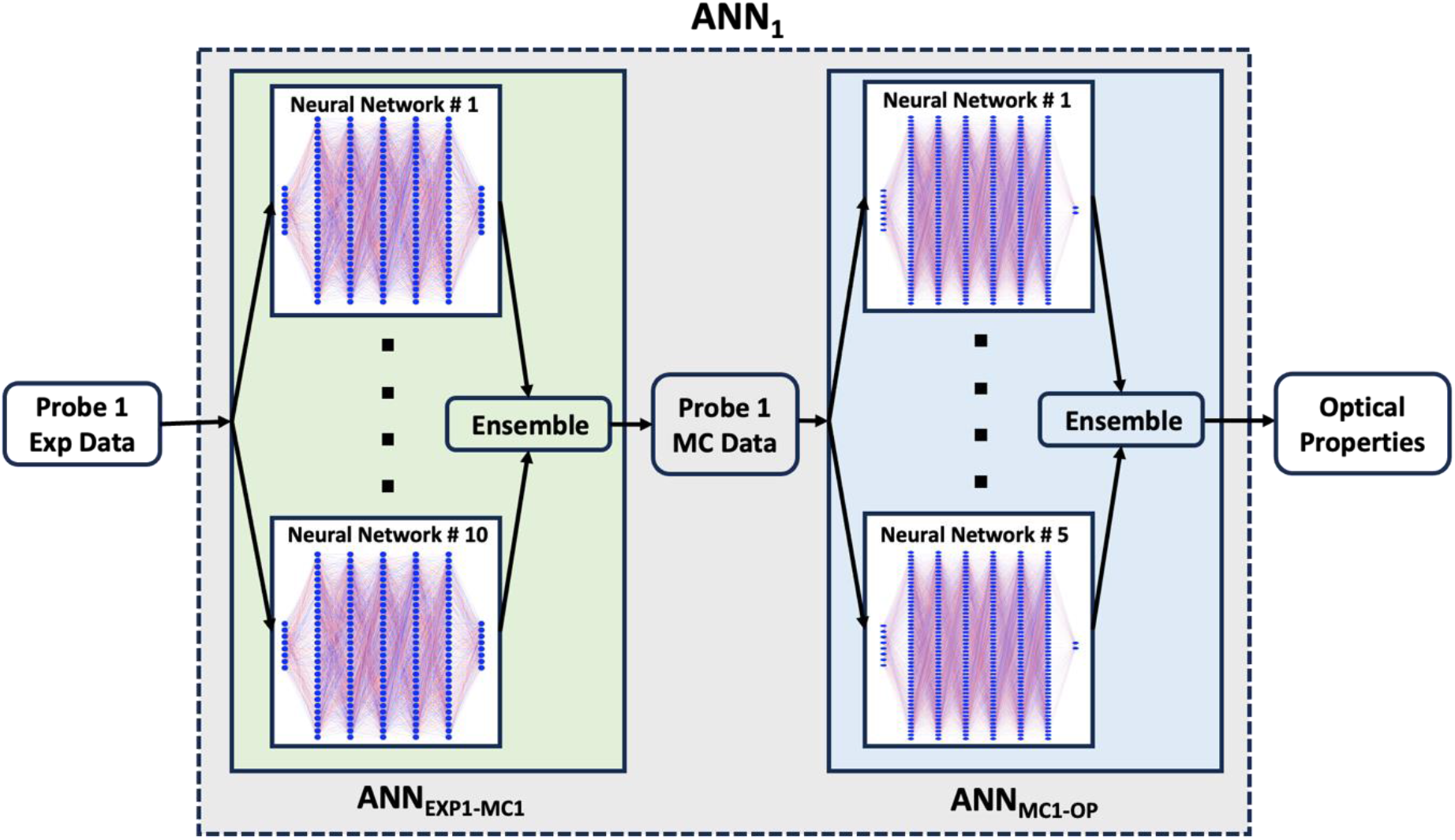
Initial ANN (ANN1) for optical property extraction, which was created by combining ANNEXP1-MC1 ensemble and ANNMC1-OP ensemble.

##### 2.3.1.1 Creation and Validation of ANN_EXP1-MC1_ Network

The ANN_EXP1-MC1_ network was an ensemble of 10 individual ANNs as shown in Figure 2. Each of these ANNs were created from the same dataset (TV_EXP1-MC1_ described in section 2.5.1.1) but with different train-validation splits and weight initializations. Given the relatively limited number of data points available for training (8848 samples), we opted for an ensemble approach to construct this network, which enhanced its stability and robustness. The input signals (experimental reflectance intensity) were not scaled. However, as the outputs (simulated reflectance intensity) of different fibers were of vastly different magnitudes, they were rescaled to [0,1] range using min-max normalization.

To train each of these ANNs, the dataset was randomly divided into two categories: Training (80%) and Validation (20%). Each individual ANN had 5 hidden layers (30 neurons in each layer) and an output layer consisting of 8 neurons resulting in 4238 parameters. The hidden layers used rectified linear unit (ReLU) activation, while the output layer employed a linear activation function. The optimal number of hidden layers and number of neurons in these layers was found using KerasTuner[15], the details of which are provided in the supplemental material. Training was continued for up to 1000 epochs. An early stopping callback was implemented to halt training if there was no improvement in the validation loss for 50 epochs, preventing overfitting. The model that demonstrated best performance on the validation set during training was restored and saved. The Adam optimizer with a mean squared error (MSE) cost function was used with a learning rate of 0.0005.

##### 2.3.1.2 Creation and Validation of ANN_MC1-OP_ Network

The optimal ANN_MC1-OP_ network was an ensemble of 5 ANNs, each with 6 hidden layers depicted as depicted in Figure 2. The details of the optimization are provided in the supplemental material. The TV_MC1-OP1_ dataset described in section 2.5.2 was used to train this network. Both the input and output data were scaled to [0,1] range. The dataset was randomly divided into training (80%) and test set (20%). An automated algorithm called Deep jointly informed neural networks (DJINN)[16] was used for automatic selection of the best ANN structure and hyperparameters without the need for time-consuming manual searching. DJINN automatically determined the near-optimal hyperparameters for our dataset, which were 250 epochs, batch size of 4916, and a learning rate of 0.007. By automating hyperparameter tuning and network structure selection, DJINN enabled us to obtain a high-performance model more efficiently. DJINN employed the Adam optimizer to minimize the mean squared error loss function (MSE). Each hidden layer utilized the rectified linear unit (ReLU) activation function, while linear activation was applied to the output layers.

Finally, ANN_EXP1-MC1_ and ANN_MC1-OP_ were integrated end-to-end to form the initial ANN (ANN_1_) as illustrated in Figure 2. To independently assess its performance, we subjected ANN_1_ to evaluation using a distinct test dataset (E_EXP1-OP1_) that was not included in training or validation. The performance of ANN_1_ was evaluated based on its ability to accurately predict both absorption (μ_a_) and reduced scattering (μ_s_′) coefficients. Predicted absorption spectra were fitted with the known chromophore basis function to retrieve absorber concentration. Predicted reduced scattering coefficient was fitted with a power-law relationship of the form μ^′^_s_ = *a (λ/λ_0_)^-b^*, where *a* and *b* are fitting coefficients, *λ* corresponds to wavelength in nm, and *λ_0_* is a normalization wavelength. This ANN_1_ served as the basis for the transfer learning technique discussed in the next section.

### 2.4 Transfer Learning

In this study, a feature extraction technique was used for transfer learning (TL). For this technique, a pre-trained model’s intermediate layers are used as feature extractors to capture relevant patterns and representations from data, followed by training additional task-specific layers on top to adapt the model for a specific task. To achieve this, only the output layer or the top few layers of the pre-trained model are made trainable while all other layers remain frozen. By training these unfrozen layers using a small transfer learning dataset, the pre-trained model is adapted to the target task. This approach is most suitable when the target dataset is similar to the data the model was pre-trained on.

Here, the ANN_EXP1-MC1_ network of ANN_1_ was used as the pre-trained model. To accomplish probe-to-probe transfer learning, only the output layer of the ANN_EXP1-MC1_ ensemble was modified. The output layer of each individual ANN of the ANN_EXP1-MC1_ ensemble was retrained using a small TL dataset collected by the target probe to create an ANN_TL_ ensemble for the target probe as shown in Figure 3. The inputs for this TL dataset were experimental reflectance signals collected from selected TL phantoms with known optical properties described in section 2.5.1, and the outputs were simulated reflectance for probe-1 obtained from the probe 1 MC library (MC_1_) described in section 2.5.2 for these known optical properties. Hence, the ANN_TL_ ensemble essentially learned the mapping from the target probe’s experimental reflectance signal to the corresponding simulated reflectance signal for probe-1. This allowed us to reuse ANN_MC1-OP_, which eliminated the need for time-consuming generation of the MC library and subsequent ANN creation for the target probe. ANN_TL_ was joined with ANN_MC1-OP_ described earlier to create an ANN_target(probe)_ that took the target probe’s experimental reflectance intensities as inputs and outputs corresponding optical properties.

**Figure 3:**
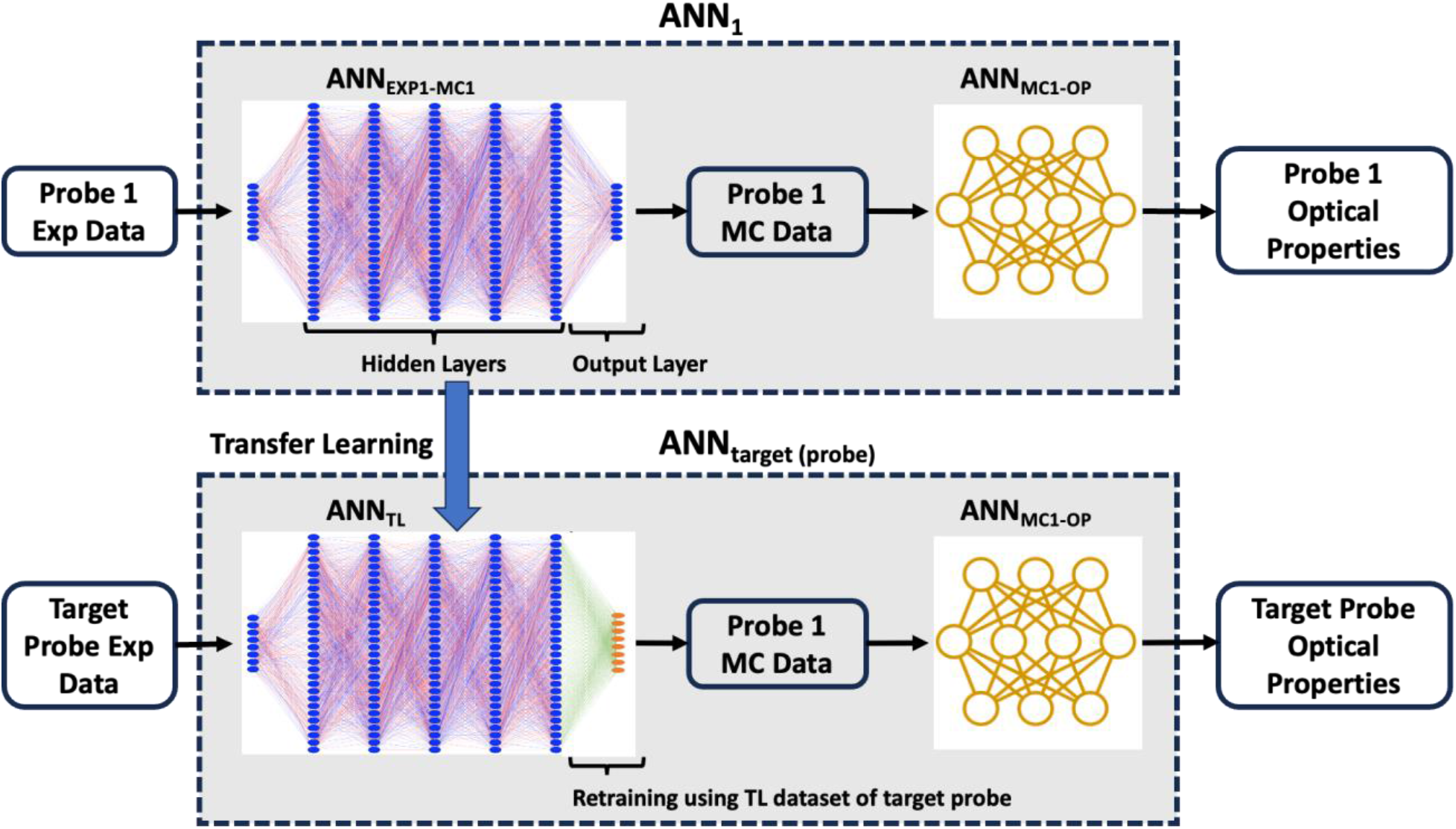
Transfer Learning to create ANNtarget (probe) from ANN1 by retraining the output layer of ANNEXP1-MC1. Although the TL algorithm was applied to all ANN of the ANNEXP1-MC1 ensemble, only one ANN is shown here for clarity.

Successful transfer learning relies on an effective transfer learning strategy, which includes the size and diversity of the transfer learning dataset, and optimal selection of hyperparameters (e.g., learning rate, epochs, batch size etc.). In order to assess our TL approach, we selected the performance of ANN_1_ on the test dataset (E_EXP1-OP1_) as the benchmark. The effective validation of our TL model ANN_target_ was determined by its capability to attain performance levels on par with those of ANN_1_. Root mean squared error (RMSE) and Mean absolute percent error (MAPE) were used as the error metrics.

In our clinical context, it is impractical to collect a large number of calibration spectra, even though having a larger number of calibration spectra with diverse conditions in the TL dataset could be beneficial. Hence, to understand the effect of TL dataset size and its diversity on the TL performance, our TL technique was applied to the pre-trained model (ANN_EXP1-MC1_) with varying sized probe-2 TL datasets. Different subsets of the TL_EXP2-MC1_ dataset were used to create different TL models and their performance were evaluated on a test dataset (E_EXP2-MC1_). This enabled us to determine which spectra should be incorporated into the TL dataset to ensure its effectiveness. In total, the TL_EXP2-MC1_ dataset consisted of 30 spectra. At first, we used the full TL_EXP2-MC1_ dataset to create the TL model. Subsequently, we reduced the size of the TL_EXP2-MC1_ dataset so that it only included phantoms with the lowest (μ_a_ = 0.05 cm^-1^) and highest absorption (μ_a_ = 1 cm^-1^) coefficient at 665 nm for each scattering condition, which resulted in 10 spectra. These spectra held particular importance for effective TL, as they spanned the outermost boundaries of the dataset. Subsequently, we used different smaller combinations of these 10 spectra for TL, down to a single spectrum.

During TL training, we randomly partitioned the TL dataset into two distinct sets: a training subset (80%) and a validation subset (20%). This validation dataset was used to prevent our TL model from overfitting to the TL dataset by using an early stopping callback (patience = 10)[15]. This low patience early stopping callback, coupled with a low learning rate (10^-4^), prevented the TL model from experiencing “catastrophic forgetting”[17] which occurs when a network forgets previously learned information when adapting to new tasks, ensuring the preservation of knowledge learned from initial probe dataset. Although the maximum number of epochs was set to 500, the training process was terminated well before reaching this limit due to the early stopping callback.

The knowledge learned from this step allowed us to create a TL algorithm that included the required number and conditions of TL spectra, and optimal hyperparameter settings for successful probe-to-probe TL. To validate the generalizability and robustness of this TL algorithm, the exact algorithm was applied to a new probe (probe-3). The TL model was created using the TL_EXP3-MC1_ dataset and performance of the resulting TL model was evaluated on an independent test dataset (E_EXP3-OP3_).

### 2.5 Datasets

#### 2.5.1 Phantom Experiments and Associated Datasets

Three stages of phantom data were collected using three different probes. In all phantoms, methylene blue (Akorn, Inc., Lake Forest, IL) was used as the absorber and Intralipid 20% (Fresenius Kabi AG, Bad Homburg, Germany) was used as the scatterer. Appropriate amounts of MB and Intralipid were mixed with 300 mL of distilled water (DI) in a black spray-painted container (Rust-Oleum Flat Black, Vernon Hills, IL) to create the phantoms described in subsections 2.5.1.1, 2.5.1.2 and 2.5.1.3. Concentration ranges of each component were selected to encompass the optical property range pertinent to our clinical application[4, 5]. Separate MB and Intralipid stocks were used for data collection for each separate probe, with each stock characterized as described below.

In order to calculate the necessary volumes of Intralipid and MB to add to DI water to achieve the desired optical properties, absorption (µ_a_) spectra of MB, scattering (µ_s_) spectra of Intralipid, and scattering anisotropy coefficients (g) for Intralipid were required. The first two were measured using a commercial spectrophotometer (Cary 50 Varian, Santa Clara, CA). To determine the scattering anisotropy of each Intralipid stock, we created 30 phantoms with μ_s_ = 25, 37.5, 62.5, 75 and 125 cm^−1^ at 665 nm, each of which had μ_a_ = 0.05, 0.1, 0.25, 0.5, 0.75, and 1.0 cm^−1^ at 665 nm from each stock. These phantoms were distinct from those used for training and/or validation. An iterative fitting algorithm was used to determine *g* by minimizing the difference between experimental signal and corresponding simulated signal obtained from the MC library described in section 2.5.2. The calculated *g* values were similar to literature values[18, 19] and showed small inter-stock variation. This small variation was relevant as it considerably affected the quality of fitting of the algorithm. This was expected because of the relatively small SDS present in our probes, making them considerably more sensitive to variations in scattering anisotropy. Hence, it was important for us to determine the value of *g* for each Intralipid stock rather than relying on the fixed values from the literature.

Our experimental phantom data collection process is briefly described here, the details of which can be found in our earlier work[4]. We first allowed the source lamp to reach a steady state over approximately 30 minutes. Calibration measurements were taken using a 3-inch calibration sphere to account for fiber throughput and lamp power variation. The probe was then placed in contact with the liquid phantom while continuously stirring. Light was emitted into the phantom using the source fiber, and diffuse reflectance spectra were collected through each of the detector fibers. For both the calibration measurements and phantom measurements, corresponding dark measurements were taken and subtracted. Then, the dark corrected spectra were normalized by the corresponding dark corrected calibration data for each detection fiber to obtain the final DRS. Each spectrum consisted of 316 wavelengths (500-740 nm).

Sub-sections 2.5.1.1, 2.5.1.2, and 2.5.1.3 detail phantom datasets and their optical properties where TV, TL, and E denote training and validation, transfer learning, and testing datasets, respectively. Figure 4 visually represents these datasets with blue, green, and orange denoting training, transfer learning, and test sets, respectively. The subscript associated each dataset is divided into two parts, with the first part indicating the input and the second part indicating the output of each dataset. EXP, MC, and OP respectively denote experimental reflectance intensity, simulated reflectance intensity, and optical properties. The numerical value in the subscript specifies the probe or MC library utilized in generating the dataset.

**Figure 4:**
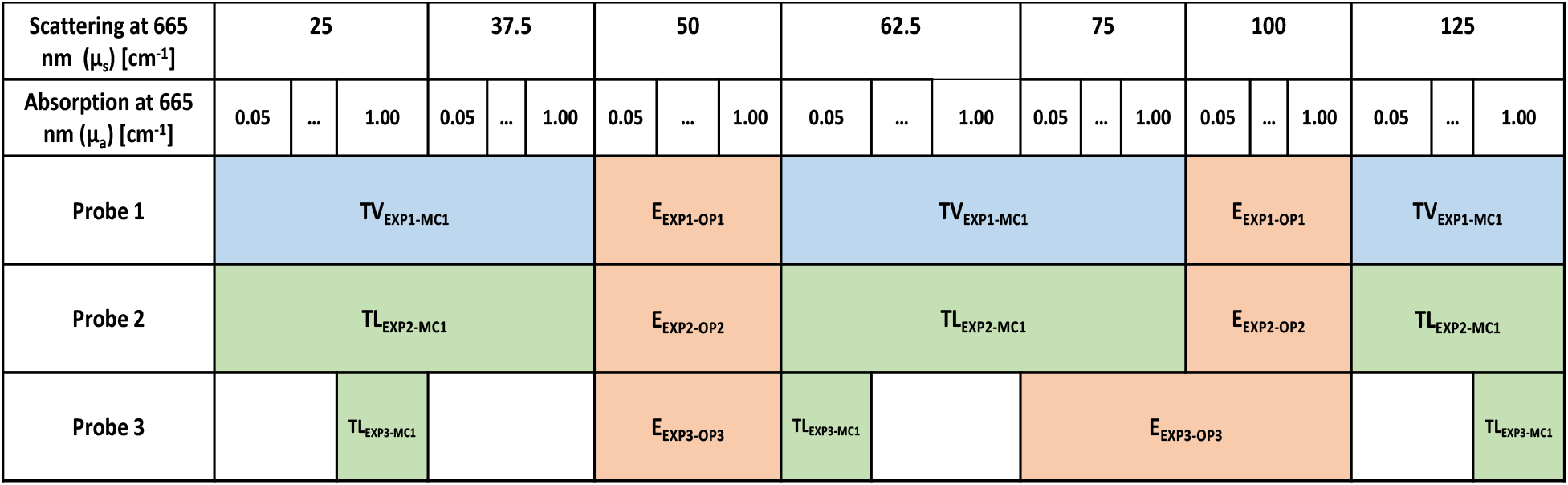
Data for different probes and their corresponding optical property ranges. The blue, green, and orange colors correspond to training, transfer learning, and test datasets, respectively.

##### 2.5.1.1 Probe-1 Dataset

Phantom data collected using probe-1 were used to create the two following datasets (see Figure 4):

- TV_EXP1-MC1_ consisted of phantoms with μ_s_ = 25, 37.5, 62.5, 75 and 125 cm^−1^ at 665 nm, each of which had μ_a_ = 0.05, 0.1, 0.25, 0.5, 0.75, and 1.0 cm^−1^ at 665 nm (30 phantoms)
- E_EXP1-OP1_ included phantoms with μ_s_ = 50 and 100 cm^−1^ at 665 nm, each of which had μ_a_ = 0.05, 0.1, 0.25, 0.5, 0.75, and 1.0 cm^−1^ at 665 nm (12 phantoms)

##### 2.5.1.2 Probe-2 Dataset

Phantom data collected using probe-2 were used to create the two following datasets (see Figure 4):

- TL_EXP2-MC1_ consisted of phantoms with μ_s_ = 25, 37.5, 62.5, 75 and 125 cm^−1^ at 665 nm, each of which had μ_a_ = 0.05, 0.1, 0.25, 0.5, 0.75, and 1.0 cm^−1^ at 665 nm (30 phantoms)
- E_EXP2-MC1_ and E_EXP2-OP2_ included phantoms with μ_s_ = 50 and 100 cm^−1^ at 665 nm, each of which had μ_a_ = 0.05, 0.1, 0.25, 0.5, 0.75, and 1.0 cm^−1^ at 665 nm (12 phantoms)

##### 2.5.1.3 Probe-3 Dataset

Phantom data were collected using probe-3 to create the following two datasets (see Figure 4):

- TL_EXP3-MC1_ comprised of the phantoms which had μ_s_ = 25 cm^−1^ and μ_a_ = 1 cm^−1^, μ_s_ = 62.5 cm^−1^ and μ_a_ = 0.05 cm^−1^, and μ_s_ = 125 cm^−1^ and μ_a_ = 1 cm^−1^. All coefficients are given for 665 nm. (3 phantoms)
- E_EXP3-OP3_ consisted of phantoms with μ_s_ = 50, 75 and 100 cm^−1^ at 665 nm, each of which had μ_a_ = 0.05, 0.1, 0.25, 0.5, 0.75, and 1.0 cm^−1^ at 665 nm (18 phantoms)

#### 2.5.2 Monte Carlo Dataset

A Monte Carlo (MC) model for probe-1 was constructed for optical property retrieval from diffuse reflectance spectra. A detailed account is of this is available in our previous publication[4]. In summary, we generated the MC library (MC_1_) by conducting graphics processing unit (GPU) accelerated MC simulations across a wide range of parameter combinations, encompassing values for absorption coefficient (μ_a_) from 0.0001 to 25 cm⁻¹ and scattering coefficient (μ_s_) from 1 to 250 cm⁻¹. These simulations were carried out using a refractive index of n=1.37 and employed the Henyey-Greenstein phase function with a scattering anisotropy factor (g) of 0.7, resulting in reduced scattering coefficients (μ_s_′) from 0.3 to 75 cm⁻¹ for the MC library. The simulations were executed using 10^8^ photon packets. The computations were performed on a Quadro RTX6000 GPU equipped with 24 GB of GPU memory (NVIDIA Corporation, Santa Clara, CA), which took about 7 days to run all optical property combinations. This MC library was used to create a TV_MC1-OP1_ dataset where the inputs were simulated reflectance intensity at 8 fibers and outputs were corresponding absorption coefficients (μ_a_) and reduced scattering coefficients (μ_s_′).

### 2.6 Statistical Analysis

Measured values are summarized as mean ± standard deviation. Paired comparisons were performed using the Wilcoxon matched-pairs signed rank test. Comparisons between the number of spectra in the transfer learning dataset used the Kruskal-Wallis test, with Dunn’s test for multiple comparisons. GraphPad Prism (v10, GraphPad Software, Inc., Boston, MA) was used for all statistical analysis.

## 3 Results

### 3.1 Performance of Initial Neural Network (ANN_1_)

#### 3.1.1 Training and Performance of ANN_EXP1-MC1_ Ensemble

Figure 5 demonstrates the progression of both training and validation losses (MSE) against the number of epochs for a representative ANN (ANN #1) selected from the ANN_EXP1-MC1_ ensemble. The plot illustrates the convergence of the ANN as training proceeded, with training and validation losses closely aligning. Early stopping callback halted the training at 544 epochs.

**Figure 5:**
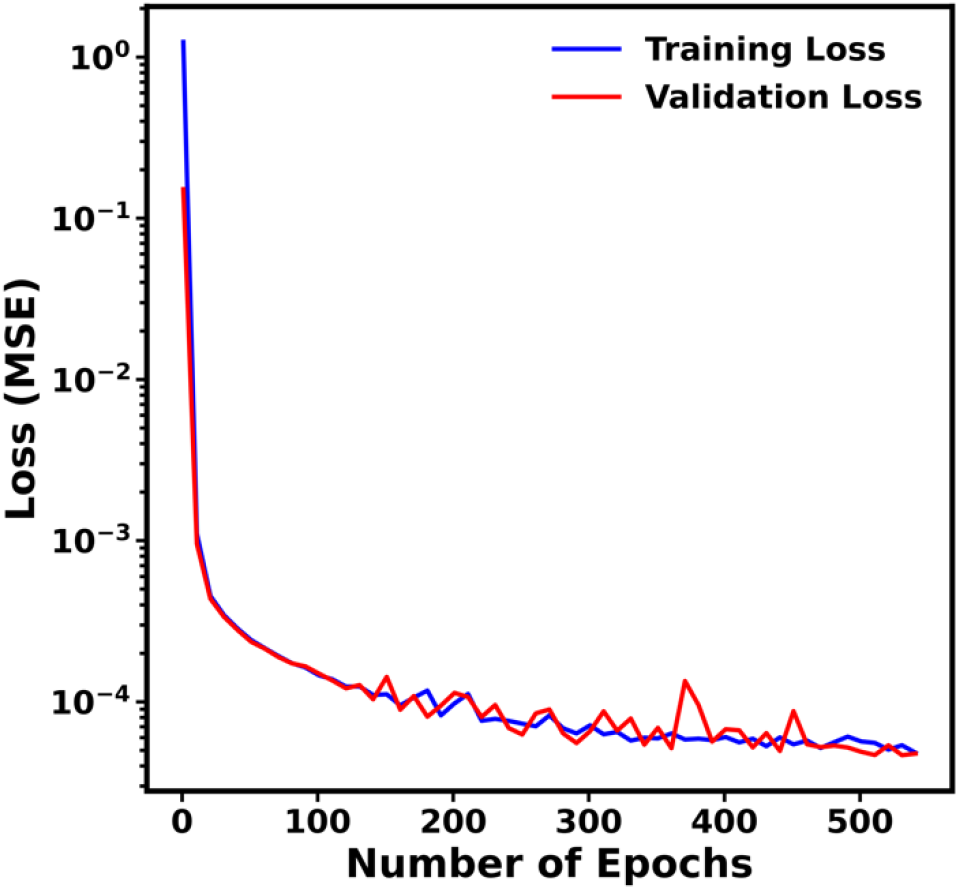
Loss (MSE) vs epochs for ANN #1 of the ANNEXP1-MC1 ensemble. Blue and red colors correspond to training and validation loss, respectively.

The performance of the ANN_EXP1-MC1_ ensemble was evaluated on the entire TV_EXP1-MC1_ dataset to ensure it properly captured the relationship between experimental and simulated reflectance signals across all detector fibers. Figure 6 demonstrates that the ensemble accurately learned the relationship between input and outputs. The RMSE and MAPE values were small across all fibers, as detailed in Table 2. The ensemble’s performance was evaluated on the entire TV_EXP1-MC1_ dataset.

**Figure 6:**
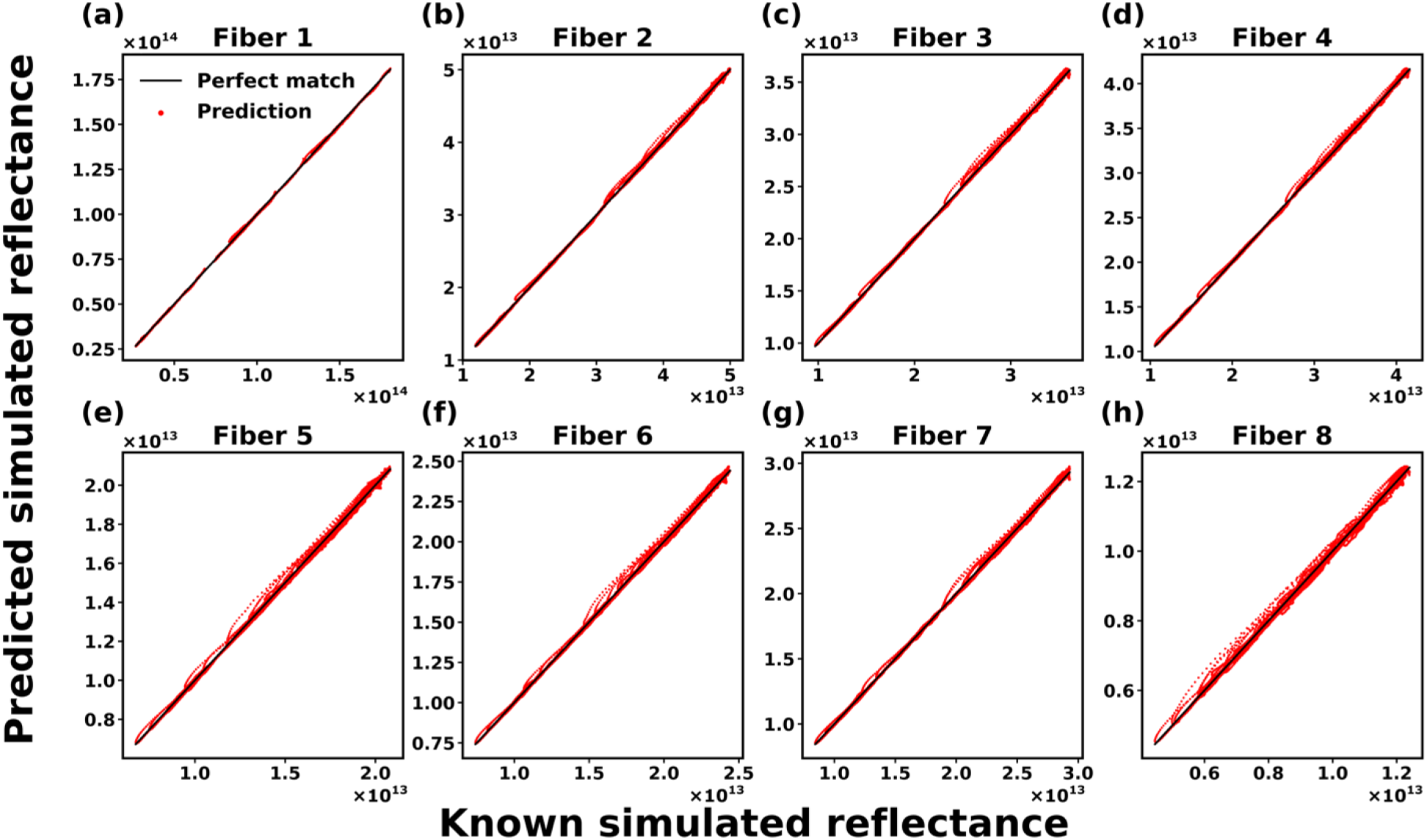
Performance of ANNEXP1-MC1 for all eight detector fibers are shown in sub-figures (a) to (h), respectively. Red dots indicate predictions and solid black lines denote perfect prediction. The units for simulated reflectance signals are detected photon weight.

**Table 2:** RMSE and MAPE for predicted simulated reflectance signal across 8 detector fibers.

| Metric | Detector Fiber 1 | Detector Fiber 2 | Detector Fiber 3 | Detector Fiber 4 | Detector Fiber 5 | Detector Fiber 6 | Detector Fiber 7 | Detector Fiber 8 |
| --- | --- | --- | --- | --- | --- | --- | --- | --- |
| RMSE | 5.34<br>$\times 10^{11}$ | 2.18<br>$\times 10^{11}$ | 1.80<br>$\times 10^{11}$ | 1.99<br>$\times 10^{11}$ | 1.36<br>$\times 10^{11}$ | 1.45<br>$\times 10^{11}$ | 1.59<br>$\times 10^{11}$ | 1.05<br>$\times 10^{11}$ |
| MAPE | 0.59% | 0.536% | 0.55% | 0.55% | 0.62% | 0.59% | 0.57% | 0.85% |

#### 3.1.2 Training and Performance of ANN_MC1-OP_ Ensemble

Figure 7 shows the training loss (MSE) vs number of epochs for the ANN_MC1-OP_ ensemble, demonstrating training convergence.

**Figure 7:**
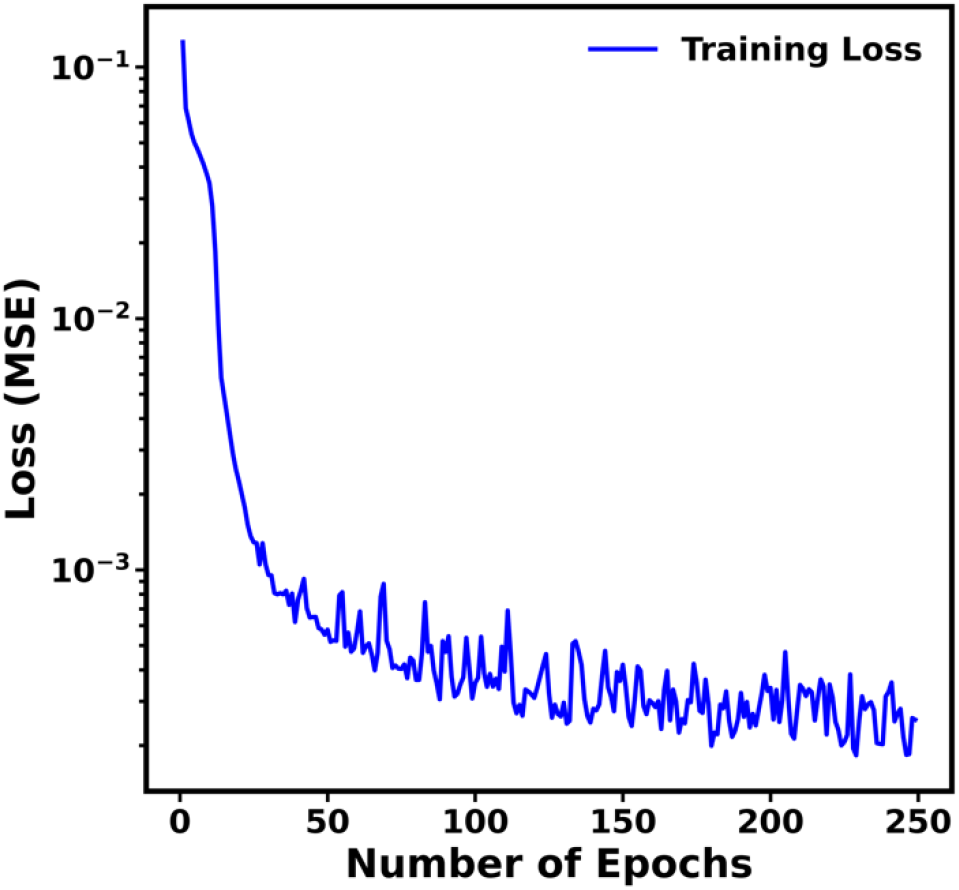
Loss (MSE) vs epoch for of ANNMC1-OP ensemble. Blue color corresponds to training loss.

The performance of the ANN_MC1-OP_ ensemble on its test set is depicted in Figure 8. Absorption coefficient (μ_a_) was predicted with a small RMSE = 0.087 cm^-1^ (MAPE = 290%). This elevated MAPE can be attributed to discrepancies arising from very small absorption coefficients (μ_a_ < 0.01 cm⁻¹). For μ_a_ > 0.01 cm^-1^, RMSE was 0.091 cm^-1^ (MAPE = 6.4%). Reduced scattering coefficient (µ_s_’) was predicted with a small RMSE = 0.48 cm^-1^ (MAPE = 1.1%). These low RMSE values indicate that ANN_MC1-OP_ successfully learned the relationship between simulated reflectance intensity and corresponding optical properties.

**Figure 8:**
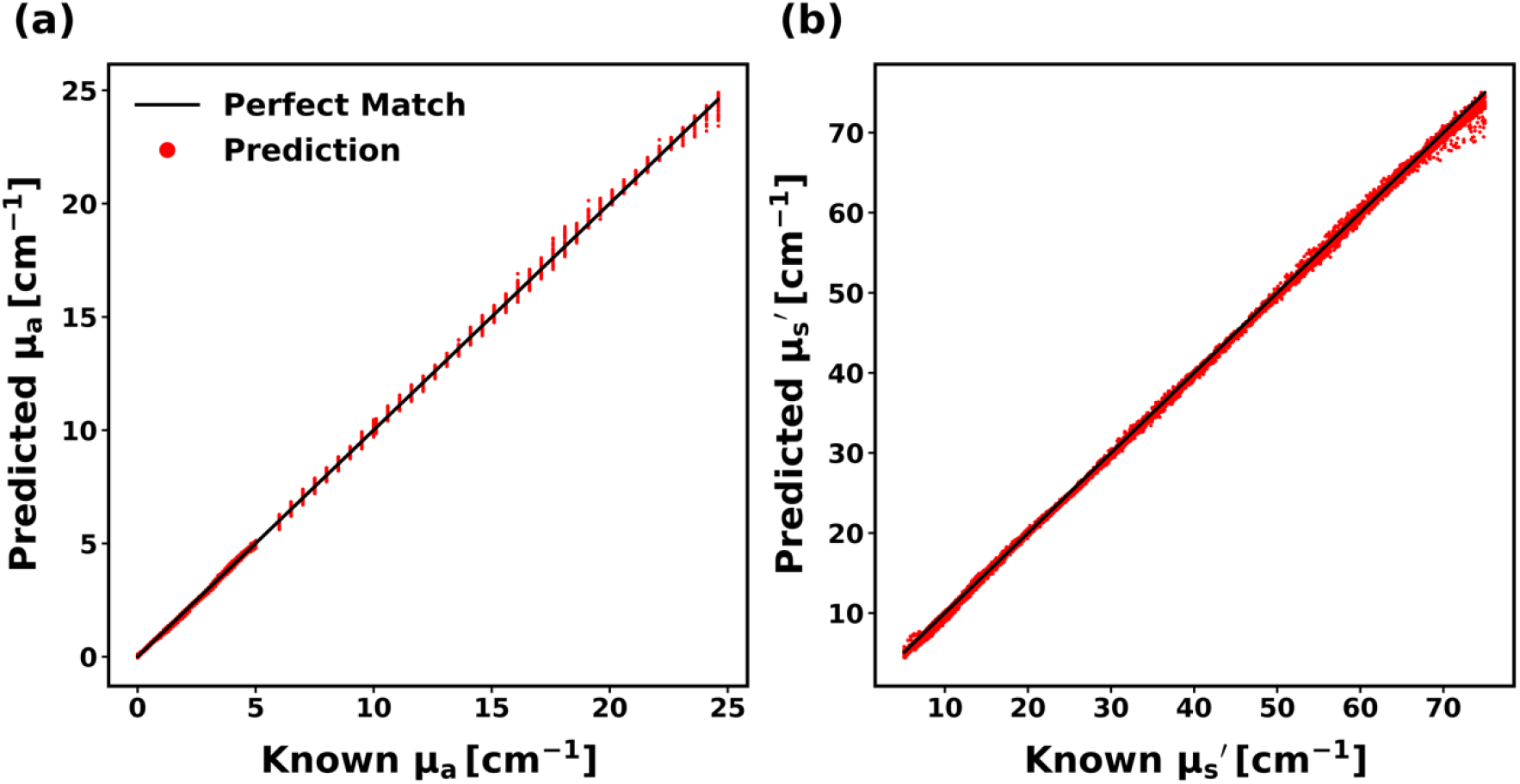
Predicted (a) absorption coefficients (μa) and (b) reduced scattering coefficients (µs’). Red dots represent predictions and solid black lines denote perfect prediction.

#### 3.1.3 Performance of the Initial ANN (ANN_1_)

The performance of the initial ANN (ANN_1_) on the test set (E_EXP1-OP1_) is presented in Figure 9. The absorption and reduced scattering spectra predictions from ANN_1_ for a representative phantom (with μ_s_ = 100 cm⁻¹ at 665 nm and μ_a_ values of 0.05, 0.1, 0.25, 0.5, 0.75, and 1.0 cm⁻¹ at 665 nm) from the test dataset are illustrated in Figures 9(a) and (b). The predicted absorption and reduced scattering spectra were fitted using the procedure mentioned in section 2.3.1.2, resulting in highly accurate fitting.

**Figure 9:**
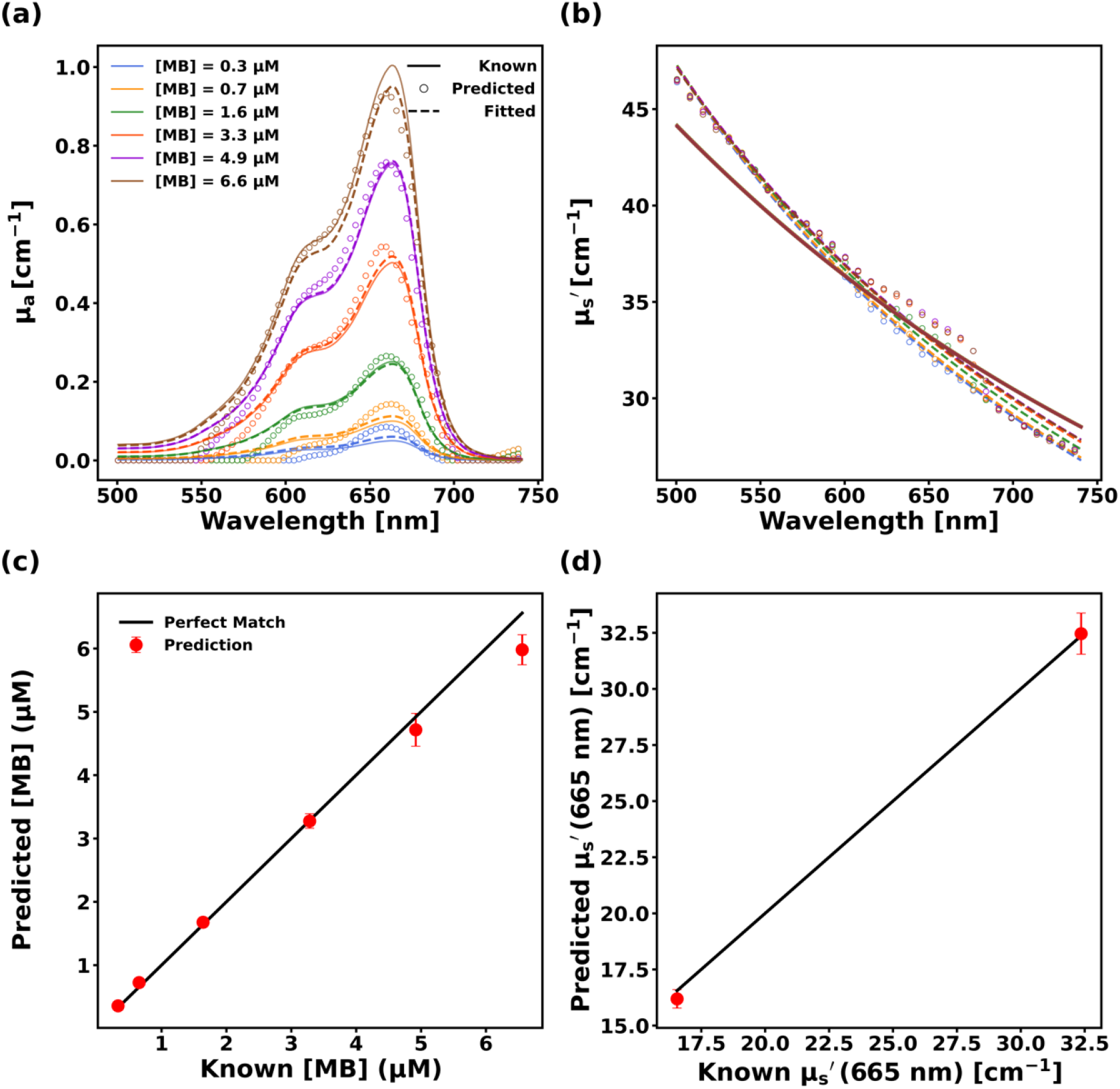
Predicted (a) absorption spectra and (b) reduced scattering spectra for a representative phantom. Solid lines, open circles, and dashed lines correspond to known, predicted, and fitted spectra respectively, while colors indicate MB concentration. Predicted (c) MB concentration and (d) µs’ at 665 nm across phantoms. Predictions are denoted by red dots and solid black lines denote perfect prediction. Symbols indicate mean recovered values across all phantom measurements at that value, while error bars correspond to standard deviation across phantoms with identical optical properties. Error bars are included in all cases but are not always visible.

Figure 9(c) illustrates that ANN_1_ accurately predicted the MB concentration of the phantoms in the test dataset with a small RMSE = 0.29 μM (MAPE = 7.46%). It also performed well in predicting the reduced scattering coefficient (µ_s_’) across all wavelengths, as evidenced by a low RMSE = 1.08 cm⁻¹ (MAPE = 3.6%). The reduced scattering coefficient at 665 nm (µ_s,665_’) was predicted with RMSE = 0.77 cm⁻¹ (MAPE = 2.54%) which is shown in Figure 9(d).

### 3.2 Transfer Learning

#### 3.2.1 Performance of Initial ANN (ANN_1_) on Probe-2 data without Transfer Learning

To access the performance of ANN_1_ for data collected with another probe, we evaluated its performance on the E_EXP2-OP2_ dataset. Figures 10(a) and (b) illustrate the absorption and reduced scattering spectra predictions of ANN_1_ for a representative phantom from E_EXP2-OP2_. ANN_1_ predicted much higher absorption than known absorption for this phantom as evident from Figure 10(a). The MB concentration and fitted reduced scattering spectra were obtained by the same methods described in sub-section 3.1.3.

**Figure 10:**
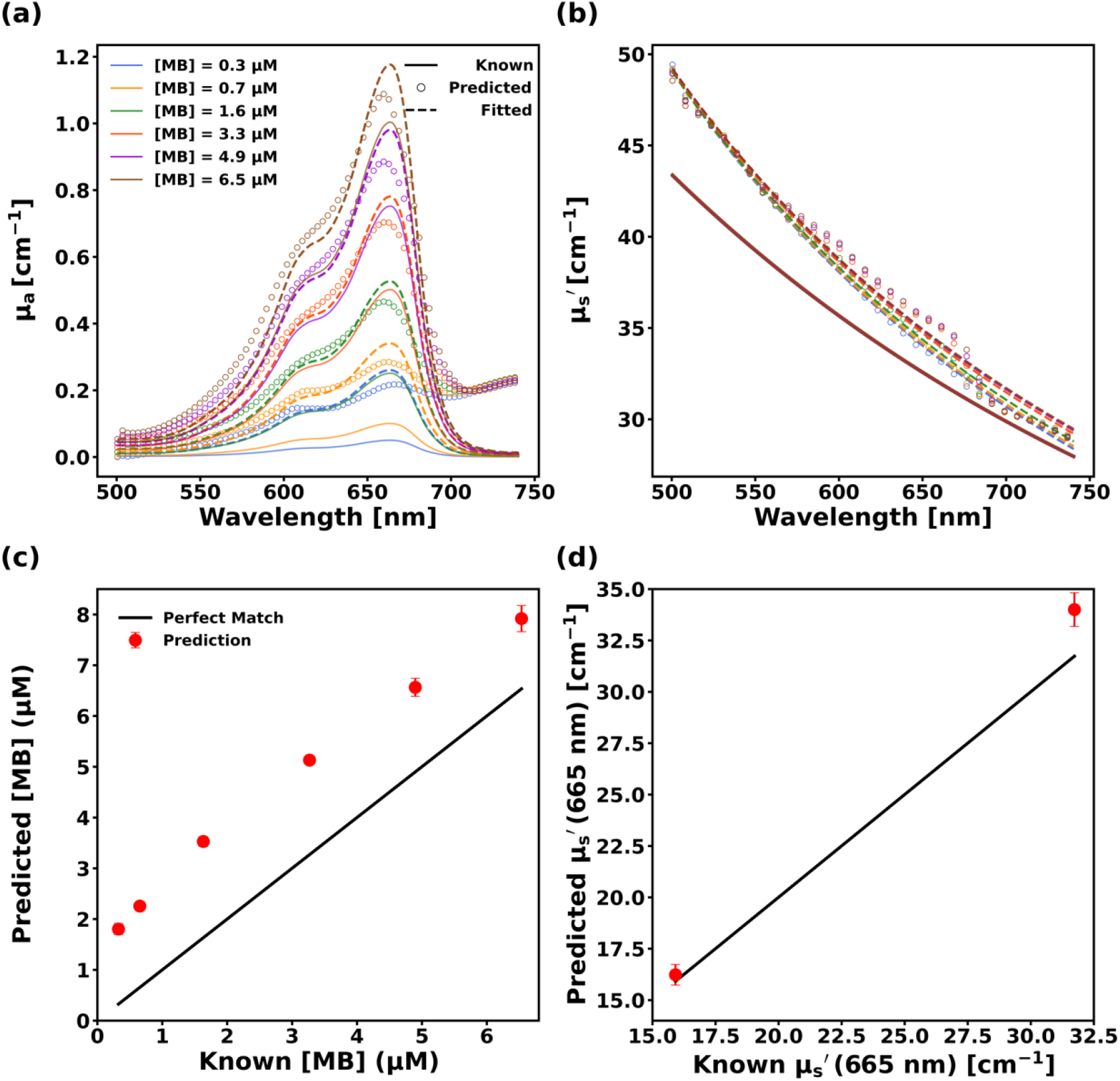
Predicted (a) absorption spectra and (b) reduced scattering spectra for a representative phantom measured with probe-2 using the neural network trained on probe-1 data (ANN1) show poor quality extraction of optical properties. Solid lines, open circles, and dashed lines correspond to known, predicted, and fitted spectra respectively while colors indicate MB concentration. Predicted (c) MB concentration and (d) µs’ at 665 nm across phantoms also show poor performance, where predictions are denoted by red dots and solid black lines denote perfect prediction. Error bars are included in all cases but are not always visible.

The failure of ANN_1_ to accurately predict MB concentration from probe-2 data can be seen in Figure 10(c), which shows the performance of ANN_1_ for the entire E_EXP2-OP2_ dataset. The MB concentration of phantoms was recovered with a large RMSE = 1.67 μM (MAPE = 154.84%). It also performed worse in predicting the reduced scattering coefficient (µ_s_’) across all wavelengths as evidenced by comparably higher RMSE = 2.2 cm⁻¹ (MAPE = 4.85%). The reduced scattering coefficient at 665 nm (µ_s,665_’) was predicted with RMSE = 1.76 cm⁻¹ (MAPE = 5.14%), as shown in Figure 10(d). These results indicate that employing ANN_1_, which was trained using probe-1 data, for optical property extraction from data collected by probe-2 results in large errors in predictions, particularly for absorption.

#### 3.2.2 Feature Extraction

From the results of sub-section 3.2.1, it is evident that usage of the initial ANN (ANN_1_) for probe-2 data leads to highly inaccurate prediction of optical properties. To adapt ANN_1_ to probe-2, we employed a feature extraction TL technique, the results of which are provided in sub-sections 3.2.2.1, 3.2.2.2, and 3.2.2.3.

##### 3.2.2.1 Number of Training Spectra vs Performance

Figure 11 presents the prediction error of different TL models created from different-sized TL training datasets on the E_EXP2-MC1_ dataset. The prediction errors of TL training datasets with the same number of spectra but varying known optical properties were used to find the mean and standard deviation of error for that particular number of spectra. Figure 11 elucidates that using 10 spectra for TL, which includes spectra with the lowest (μ_a_ = 0.05 cm^-1^) and highest absorption (μ_a_ = 1 cm^-1^) coefficient at 665 nm for each scattering condition, leads to identical error compared to that of using all 30 spectra in the TL_EXP2-MC1_ dataset (0.021 vs. 0.021). We found that as we incorporate smaller and smaller subsets of these 10 spectra in the TL dataset (from 5 spectra to 1 spectrum), the performance of the resultant TL model worsens, indicated by the higher MSE shown in Figure 11. The usage of a smaller number of spectra also leads to a higher variance in the error compared to the usage of a higher number of spectra.

**Figure 11:**
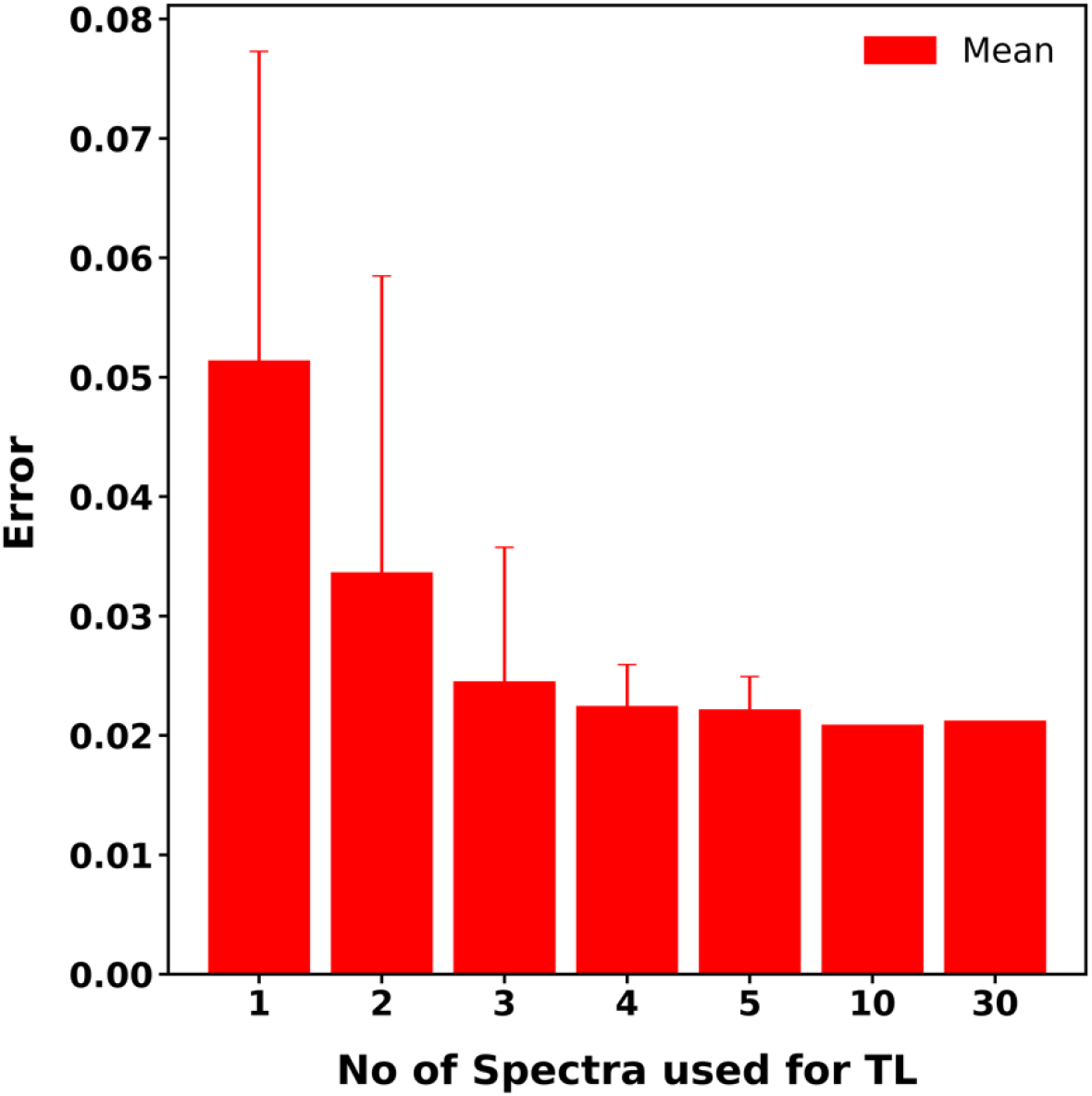
Prediction error for ANNTL models created from TL datasets with varying number of training spectra. The X-axis denotes the number of spectra in the TL dataset, and the Y-axis denotes the prediction error. Solid bars represent mean values across all combinations of the specified number of spectra, with error bars corresponding to standard deviation.

##### 3.2.2.2 Selection of Spectra and Creation of TL Algorithm

We chose to utilize 3 training spectra in our TL algorithm, as we can see that utilizing 3 spectra for TL results in comparable error to utilizing all 30 spectra (0.024 vs. 0.021). Utilization of only 2 training spectra resulted in an increase in error (0.034 vs. 0.024). To select the optimal spectra to be incorporated into the TL dataset, we considered the strengths and weaknesses of ANNs. ANNs work accurately and reliably only within the range of the data they were trained on. Beyond this range, performance can degrade quite rapidly[20]. Hence, to maximize the range of validity of our TL model, we decided to integrate two spectra from the extremes of the TL dataset (TL_EXP2-MC1_). Specifically, we included the spectrum with the lowest scattering and highest absorption (μ_s_ = 25 cm^−1^ and μ_a_ = 1 cm^−1^ at 665 nm), as well as the spectrum with the highest scattering and absorption (μ_s_ = 125 cm^−1^ and μ_a_ = 1 cm^−1^ at 665 nm). Moreover, we also included an intermediate spectrum (μ_s_ = 62.5 cm^−1^ and μ_a_ = 0.05 cm^−1^ at 665 nm) to improve interpolation. In this step, we also determined the optimal hyper-parameters for TL (e.g., learning rate, epochs, etc.). Based on these, a TL algorithm was created for probe-to-probe TL which will allow us to create a ANN_target(probe)_ for the target probe optical property recovery.

The algorithm was:

1. Collection of 3 spectra from 3 known phantoms using the target probe to create the TL dataset. The phantoms are:

a. μ_s_ = 25 cm^−1^ and μ_a_ = 1 cm^−1^ at 665 nm
b. μ_s_ = 62.5 cm^−1^ and μ_a_ = 0.05 cm^−1^ at 665 nm
c. μ_s_ = 125 cm^−1^ and μ_a_ = 1 cm^−1^ at 665 nm
2. Usage of the TL dataset to adapt ANN_EXP1-MC1_ to create ANN_TL_ adapted for target probe
3. Integration of ANN_TL_ with ANN_MC1-OP_ to create ANN_target(probe)_ which is capable of recovering optical properties from DRS collected by the target probe.

##### 3.2.2.3 Performance of ANN_target(2)_ on Probe-2 Data

This algorithm was applied to ANN_1_ to create ANN_target(2)_ for probe-2 and its performance was evaluated on E_EXP2-OP2_. Figures 12(a) and (b) illustrate the absorption and reduced scattering spectra predictions of ANN_target(2)_ for a representative phantom from E_EXP2-OP2_. We observe much better agreement between the known and predicted spectra for both absorption and reduced scattering compared to the incorrect insertion of probe-2 data into ANN_1_.

**Figure 12:**
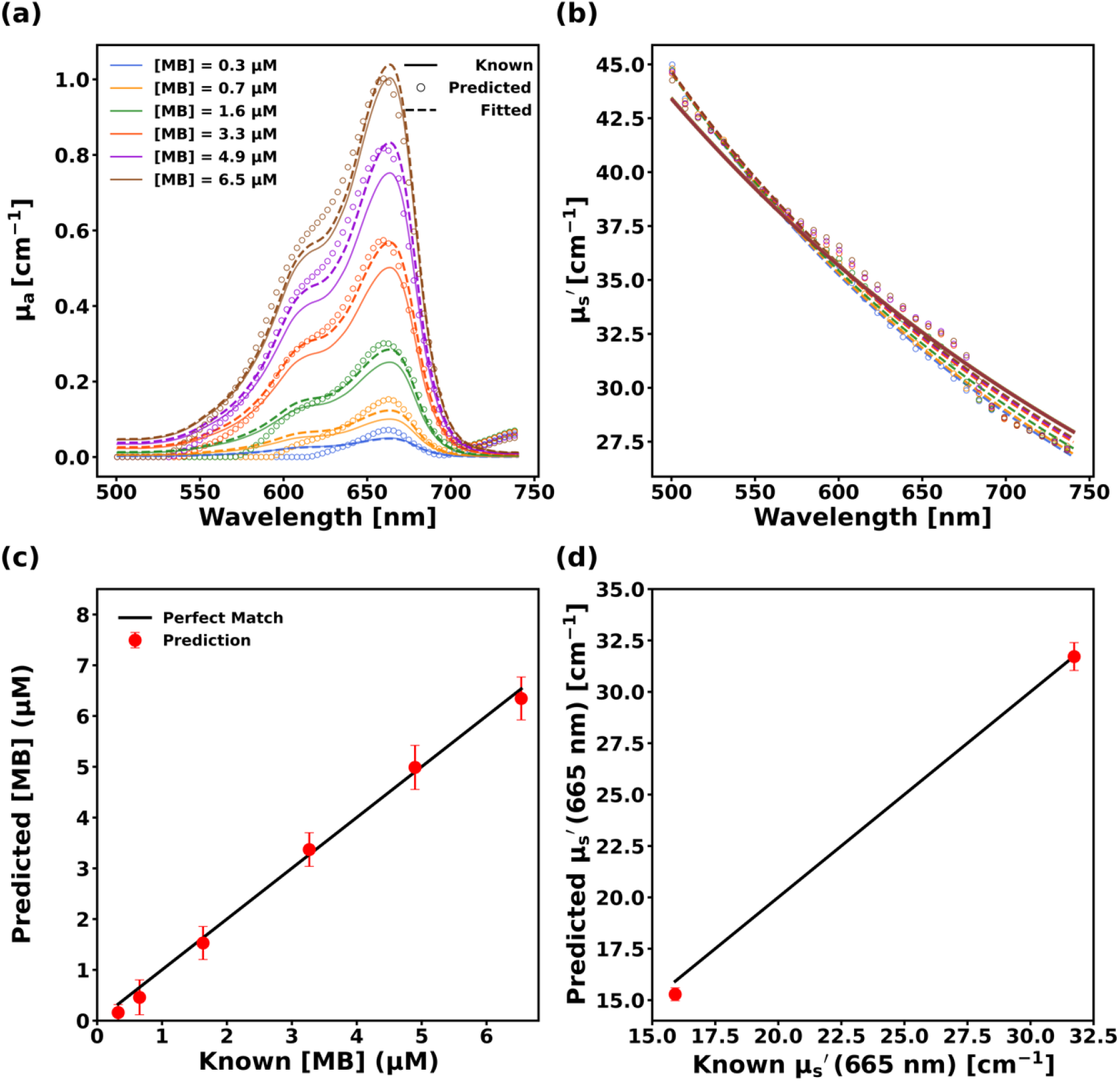
Predicted (a) absorption spectra and (b) reduced scattering spectra for a representative phantom measured with probe-2 after transfer learning. Solid lines, open circles, and dashed lines correspond to known, predicted, and fitted spectra respectively while colors indicate MB concentration. Predicted (c) MB concentration and (d) µs’ at 665 nm across phantoms, where predictions are denoted by red dots and solid black lines denote perfect prediction. Error bars are included in all cases but are not always visible.

The ability of ANN_target(2)_ to accurately predict MB concentration can be seen in Figure 12(c), which shows the performance of ANN_target_ for the entire E_EXP2-OP2_ dataset. The MB concentration of phantoms was recovered with a small RMSE = 0.38 μM (MAPE = 24.82%). It also showed superior performance in predicting the reduced scattering coefficient (µ_s_’) across all wavelengths as evidenced by low RMSE = 0.88 cm⁻¹ (MAPE = 3.68%). The reduced scattering coefficient at 665 nm (µ_s,665_’) was predicted with RMSE = 0.71 cm⁻¹ (MAPE = 3.01%) which is shown in Figure 12(d).

Overall, ANN_target(2)_ exhibited significantly improved performance on the E_EXP2-OP2_ dataset in comparison to ANN_1_ both in predicting MB concentration (p = 0.0005) and reduced scattering coefficient at 665 nm (p = 0.0005).

### 3.2.3 Application of TL Algorithm to Probe-3

The performance of ANN_1_ on probe-3 data (E_EXP3-OP3_) is shown in Figures 13(a) and (b) as the pre-TL prediction. The MB concentration prediction error was huge, with a RMSE = 4.1 μM (MAPE = 361.08%). For phantoms with μ_a_ > 0.1 cm^-1^, MB concentration was predicted with RMSE = 4.06 μM (MAPE = 124.51%). Prediction of the reduced scattering coefficient (µ_s_’) across all wavelengths was also poor, with RMSE = 2.47 cm⁻¹ (MAPE = 5.68%). The reduced scattering coefficient at 665 nm (µ_s,665_’) was predicted with an RMSE = 2.08 cm⁻¹ (MAPE = 5.45%) as shown in Figure 13(b).

**Figure 13:**
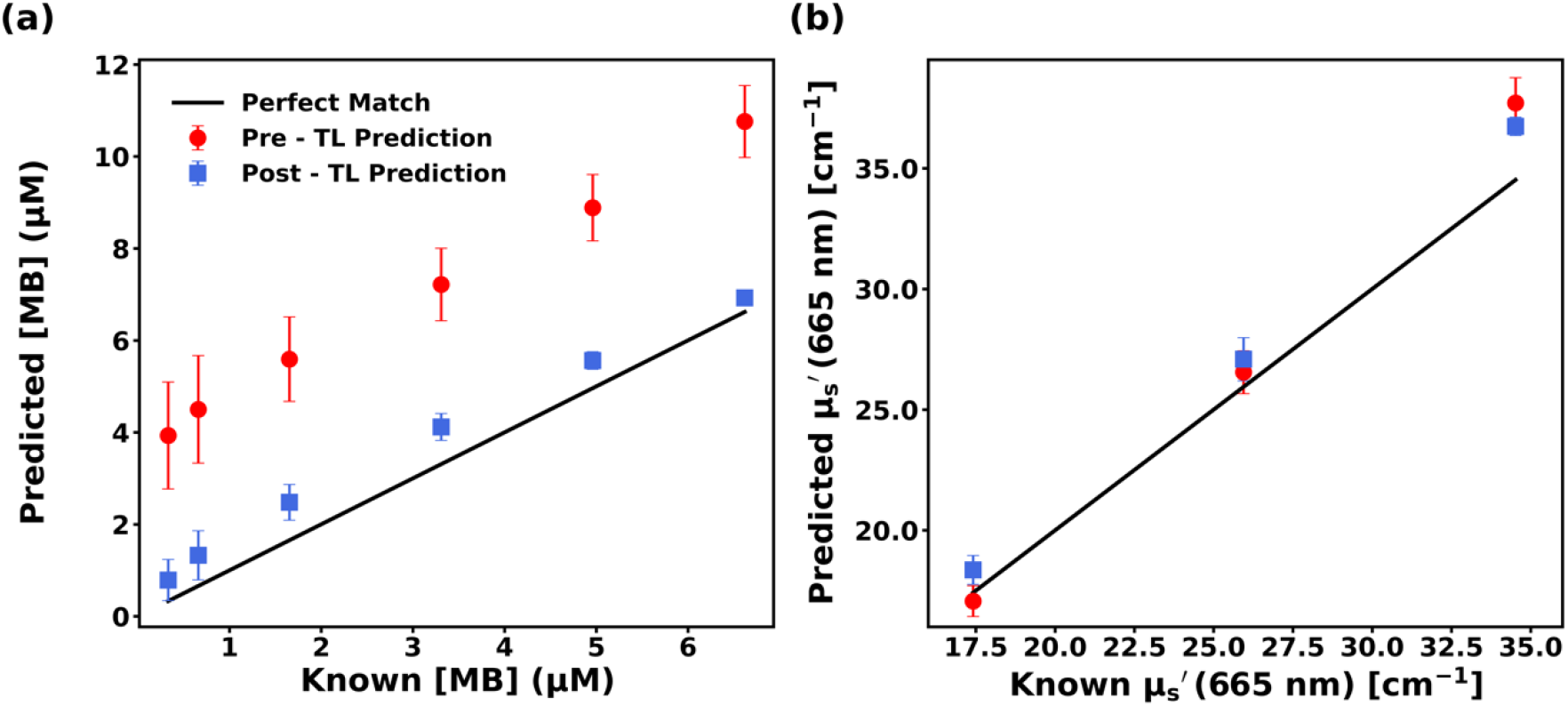
Predicted (a) MB concentration and (b) µs’ at 665 nm for phantoms measured with probe-3 for both before (pre-TL) and after (post-TL) the application of transfer learning. Pre-TL and post-TL predictions are denoted by red dots and blue squares, respectively, while solid black lines denote perfect prediction. Error bars are included in all cases but are not always visible.

The TL algorithm described in Section 3.2.2.2 was applied to ANN_1_ to create ANN_target(3)_ for probe-3 and its performance was also evaluated on E_EXP3-OP3_, which are shown in Figures 13(a) and (b) as the post-TL prediction. The MB concentration of phantoms was recovered with a RMSE = 0.73 μM (MAPE = 61.89%), as depicted in Figure 13 (a). This high MAPE can be attributed to mismatch in prediction of low-absorption phantoms for which slight mismatch in prediction causes huge MAPE values. For phantoms with μ_a_ > 0.1 cm^-1^, MB concentration was predicted with RMSE = 0.72 μM (MAPE = 22.8%). It was also able to predict reduced scattering coefficient (µ_s_’) across all wavelengths with a low RMSE = 1.67 cm⁻¹ (MAPE = 4.11%). The reduced scattering coefficient at 665 nm (µ_s,665_’) was predicted with RMSE = 1.68 cm⁻¹ (MAPE = 5.43%) which is shown in Figure 13(b). ANN_target(3)_ exhibited significantly improved performance on the E_EXP3-OP3_ dataset in comparison to ANN_1_ for MB concentration prediction (p < 0.0001), while prediction of the reduced scattering coefficient at 665 nm was not significantly improved (p = 0.2).

## 4 Discussion

In this study, we created and validated an ANN that can accurately predict optical properties from DRS collected by a specific fiber optic probe. Then, we investigated utilization of this ANN for optical property prediction from DRS collected by other probes having dissimilar SDS. We found that doing so led to large errors in prediction. To solve this issue, we utilized a feature-extraction based transfer learning algorithm. We showed that this algorithm can reliably adapt an ANN created for one probe to a target probe for accurate optical property recovery by the target probe. This probe-to-probe calibration can be achieved by using a small transfer learning dataset consisting of only three spectra collected by the target probe.

Other groups have used a similar ANN approach for optical property recovery from DRS. Ivančič *et al* predicted μ_a_ with RMSE = 0.09 cm^-1^ (relative RMSE = 2.6%) and μ_s_′ with RMSE = 0.26 cm^-1^ (relative RMSE = 1.6%) at SDS ranging from 220-1200 μm[21]. However, their performance evaluation was limited to a single low scattering and high absorption phantom. Furthermore, they employed separate ANNs for predicting each optical parameter which resulted in four distinct ANNs, whereas we used a single ANN for predicting both absorption and reduced scattering spectra. Chong *et al* demonstrated recovery of μ_a_ and µ_s_’ with mean relative error ranging from 1.62-5.92% and 0.8-1.6%, respectively, with their baseline neural network and its physics-guided variants, but only for a simulation dataset[20]. An *et al* developed ANNs for their spectroscopy system, which predicted μ_a_ with mean absolute error ranging from 0.13-0.17 cm^-1^ and μ_s_′ ranging from 1.43-1.71 cm^-1^ [22]. However, their ANN prediction was limited to only four wavelengths compared to our broad-spectrum prediction ranging from 500-740 nm. Lan *et al* predicted μ_a_ with Euclidean distance = 0.25 and μ_s_′ with Euclidean distance = 4.14, which is somewhat difficult to interpret[14]. With our initial ANN (ANN_1_), we achieved comparable performance to these previous studies. However, these prior publications only investigated the performance of ANN-based optical property recovery for a single instance of their respective fiber optic probes. The effects of SDS mismatch between probes on the recovered optical properties was not previously explored. This is vitally important for applications requiring multiple copies of a designed fiber optic probe, or single use devices.

We have shown that ANNs for optical property extraction are probe-specific. In other words, an ANN created for one probe can’t be used for another probe if their SDS are considerably different. This is particularly true for the short SDS required by our clinical fiber optic probe. Precise knowledge of SDS is therefore required for traditional methods, such as Monte Carlo lookup tables[4, 11] or analytical approximations[10], as well as generation of Monte Carlo models for ANNs trained on simulation data. Hence, each time we intend to extract optical properties using a new probe, we are required to perform time-consuming probe characterization, probe-specific MC library creation, and phantom data collection, as well as ANN creation and validation to ensure satisfactory performance with this new probe. These steps require a substantial amount of effort, technical expertise, and time (7-14 days). These factors can severely limit the clinical implementation of an ANN-based spectroscopy system for our envisioned multi-center clinical trials or to transform this system into a widely-used product on a large scale[23].

As we have demonstrated, our transfer learning algorithm offers a possible solution to these problems. It eliminates the need for performing the aforementioned lengthy and time-consuming probe-specific MC library and ANN creation for each new probe, reducing the time and expertise required. Even without code optimization, our probe-to-probe calibration can be done rapidly within 10 minutes, so it can be easily integrated into a clinical workflow. We envision that this would utilize well-characterized solid phantoms for transfer learning, which have been shown to be highly reproducible[24]. Furthermore, this transfer learning algorithm could potentially reduce the cost of probe manufacturing. As we have shown that our algorithm allows for accurate recovery of data from probes with relatively large differences in SDS from those specified, this could allow for the utilization of less expensive probes with looser tolerances on fiber positioning. This could be extended to the utilization of sterile disposable probes for straightforward integration into the clinical workflow.

We observed significantly better prediction performance after the application of the transfer learning algorithm for both probes 2 and 3. For probe-2, errors in both MB concentration and scattering coefficient prediction were reduced up to 77.2% and 60%, respectively. For probe 3, errors in both MB concentration and scattering coefficient prediction were reduced up to 82.2% and 32.4%, respectively. These findings indicate that the transfer learning algorithm enhanced the prediction accuracy of both probes 2 and 3, demonstrating the generalizability of our proposed TL algorithm.

We have shown that the magnitude of prediction error depends on the SDS mismatch between the initial probe and the target probe. This was apparent when data from probe-2 or 3 were used as inputs to the initial ANN (ANN_1_) without transfer learning. For probe-2 data, the ANN_1_ predicted MB concentration with RMSE = 1.67 μM (MAPE = 154.84%) whereas for probe 3 data, it predicted MB concentration with RMSE = 4.1 μM (MAPE = 361.08%) which is 2.45 times higher. Between probes 1 and 2, the SDS mismatch had RMSE = 39.69 μm (MAPE = 3.46%) whereas between probes 1 and 3, the SDS mismatch had RMSE = 100.91 μm (MAPE = 10.94%) which is 2.54 times higher. It is evident that a higher mismatch in SDS leads to higher prediction errors. Due to this greater mismatch between fiber positioning between probes 1 and 3 compared to that of probes 1 and 2, even after the application of transfer learning the MB concentration recovery had somewhat higher RMSE = 0.73 μM (MAPE = 61.89%) for probe 3. This implies that there may be a maximum SDS difference that can be tolerated by this transfer learning approach. Unfortunately, we do not have a sufficient selection of probes to test this directly, but this will be an area of future study.

Although our proposed transfer learning algorithm has great potential in facilitating multi-center trials and ultimately widespread adaption of probe-based optical property recovery using ANN, it is not without limitations. We employed only three probes in this study which prevents us from creating a general guideline involving tolerance in fiber positioning and its effect on optical property prediction. The differences among the probes were substantial but not overwhelmingly large. In future research, we are planning to investigate these issues more thoroughly by employing a larger number of probes with varying SDS. One weakness of the ANN approach is the inability to perform accurately and reliably outside the training range[25]. Although the ANNs used in this study were trained on datasets that covered the range pertinent to our clinical application, there is a possibility that these ANNs will encounter out-of-domain data in a clinical scenario. We plan to utilize physics-guided neural networks for optical recovery in our future endeavors as these networks have shown superior out-of-domain performance for different tasks[20]. The reliability of our TL algorithm may be compromised by uncertainties in the knowledge of the optical properties for the phantoms used for transfer learning. This could be mitigated by the usage of solid phantoms with identical optical properties, including regular validation of these phantoms with another well-validated spectroscopy system.

## Code, Data, and Materials

Data are available upon reasonable request from the authors.

## Acknowledgments

This work was funded by grant EB029921 from the National Institutes of Health.

