## Supplemental Figures for "Application of Transfer Learning for Rapid Calibration of Spatially-resolved Diffuse Reflectance Probes for Extraction of Tissue Optical Properties"

### Supplemental Materials:

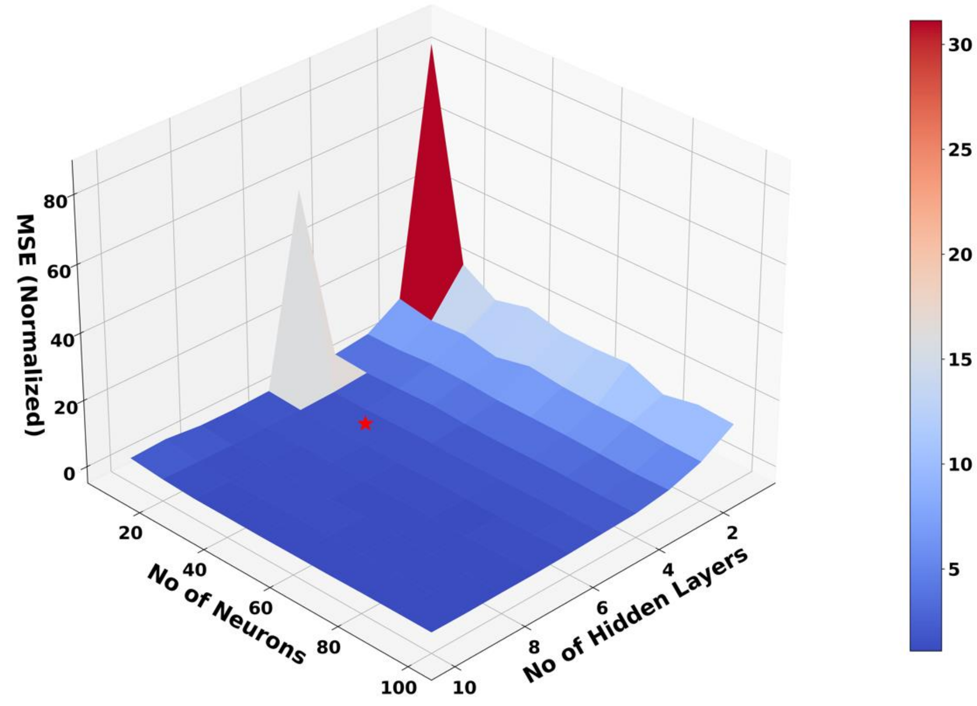


**Figure S1:** Surface plot showing the performance of different $ANN_{EXP1\text{-}MC1}$ networks on a validation dataset. The number of hidden layers were varied from 1 to 10, and the number of neurons in each hidden layer were varied from 10 to 100 in an increment of 10, resulting in 100 different combinations. The mean squared error (MSE) for this validation set is a function of the number of hidden layers and the number of neurons in each hidden layer. The displayed MSE is normalized by the smallest MSE error of all combinations for ease of visualization. The selected $ANN_{EXP1\text{-}MC1}$ network (5 hidden layers, 30 neurons in each hidden layer) is denoted with a red star.

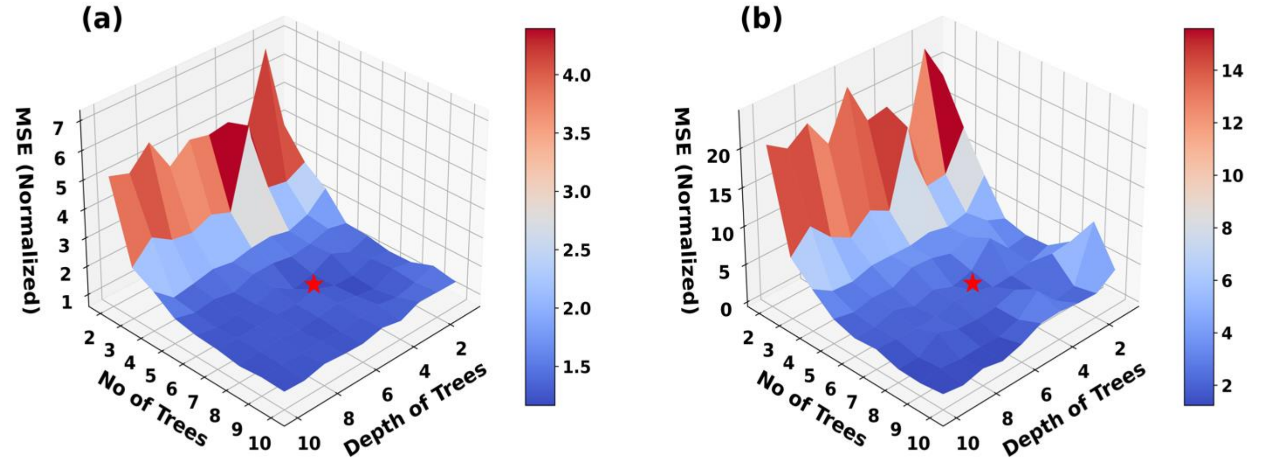


**Figure S2:** Surface plots showing the performance of different $ANN_{MC1-OP}$ networks on a validation dataset for (a) absorption coefficient and (b) reduced scattering coefficient. Both the number of trees and depth of trees were varied from 1 to 10, resulting in 100 different ANN models. The mean squared error (MSE) on this validation set is a function of the number of trees (number of ANNs) and depth of trees (number of layers). Displayed MSE for each are normalized by their respective smallest MSE for all combinations. The selected $ANN_{MC1-OP}$ network (5 trees, depth of trees = 6) is denoted with a red star.
